# Enforcing local DNA kinks by sequence-selective trisintercalating oligopeptides of a tricationic porphyrin. A polarizable Molecular Dynamics study

**DOI:** 10.1101/2023.07.26.550640

**Authors:** Nohad Gresh, Krystel El Hage, Louis Lagardère, Frédérique Brégier, Jérémy Godard, Jean-Philip Piquemal, Martine Perrée-Fauvet, Vincent Sol

## Abstract

Bisacridinyl-bisarginyl porphyrin (BABAP) is a trisintercalating derivative of a tricationic porphyrin, formerly designed and synthesized in order to selectively target and photosensitize the ten-base pair palindromic sequence d(CGGGCGCCCG)_2_. We resorted to the previously derived (Far et al., 2004) lowest energy-minimized (EM) structure of the BABAP complex with this sequence as a starting point. We performed polarizable molecular dynamics (MD) on this complex. It showed, over a 150 ns duration, the persistent binding of the Arg side-chain on each BABAP arm to the two G bases upstream from the central porphyrin intercalation site. We subsequently performed progressive shortenings of the connector chain linking the Arg-Gly backbone to the acridine, from n=6 methylenes to 4, followed by removal of the Gly backbone and further connector shortenings, from n=4 to n=1. These resulted into progressive deformations (‘kinks’) of the DNA backbone. In its most accented kinked structure, the DNA backbone was found to have a close overlap with that of DNA bound to *Cre* recombinase, with, at the level of one acridine intercalation site, negative roll and positive tilt values consistent with those experimentally found for this DNA at its own kinked dinucleotide sequence. Thus, in addition to their photosensitizing properties, some BABAP derivatives could induce sequence-selective, controlled DNA deformations, which are targets for cleavage by endonucleases or for repair enzymes.

## Introduction

The d(GGCGCC)_2_ palindrome is encountered in oncogenic (1) and retroviral sequences (2), and was recently found at three insertion sites of *Alu* repeats (3). *Alu* repeats are transposon polynucleotides which can detach from the genome and integrate into other DNA sequences, as well as into extrachromosomal circulating DNA (ec-DNA). Ec-DNA is identified as a major cause in tumor progression, and is a much more accessible target than nuclear DNA (4) (5). Such occurrences are a strong incentive to design DNA-binding ligands which could bind with augmented selectivity to this palindrome compared to all possible hexameric sequences, of which there is a total of 2080 (6). Toward this aim, we have previously designed oligopeptide derivatives of two intercalators. The first is the anthraquinone ring of the antitumor drugs ametantrone (AMT) and mitoxantrone (MTX) (7). The second is a cationic porphyrin, which acts as a photosensitizer in dynamic phototherapy (8) (9). We previously reported the results of joint computational and experimental studies regarding both anthraquinone (10) and porphyrin (11) oligopeptides. A short-duration molecular dynamics (MD) study was reported in a 2010 paper (10), but most computational studies had resorted so far to energy-minimization. Advances in the Tinker-HP software (12) recently operational on GPU supercomputers (13) have paved the way to long-duration MD simulations of the complexes of oligonucleotides with newly-designed mono- and trisintercalating MTX derivatives (14). Such simulations are now extended to bisacridinyl-bisarginyl porphyrin (BABAP) and derivatives.

The molecular structure of BABAP is represented in Figure 1. In its design, while the porphyrin ring intercalates at the central d(CpG)_2_ step of the palindrome, the Arg side-chain of each arm binds along one DNA strand in the major groove to the two successive G bases upstream. The Arg-Gly backbone is connected to an amino-acridine (AA) by a C6-paraffinic chain, and each AA ring intercalates at a d(CpG)_2_ site upstream. Thus the overall targeted sequence is d(C GGGC GCCC G)_2_, a blank denoting an intercalation site. Both circular dichroism experiments and topoisomerase I unwinding experiments lent support to trisintercalation in the BABAP complex with an oligonucleotide with this decanucleotide as a central sequence, while mono- and bisintercalation took place, respectively, with BAP (devoid of the two AA extension) and MABAP having one single AA intercalator (11). The present study starts from the lowest energy-minimum structure of the complex reported in (11), in which intercalation of both AA arms takes place from the major groove. We first evaluate the extent to which long-duration MD with a multipolar polarizable potential in a water bath with counterions retains the most important feature from EM. Polarizable potentials are known to display high accuracy for polar and charged systems, including proteins and nucleic acids (15-18). Because AA intercalation is found to take place without significant distortion of the sugar-phosphate backbone between the central and both AA intercalation sites, namely at steps d(GGGC), this led us to next address the question: as long as AA intercalation remains enabled, what could be the impact of progressive shortenings of the connector on the conformation of this step, and could local ‘kinks’ be sequence-selectively enforced by some shorter-chain BABAP derivatives?

**Figures 1a-d.**
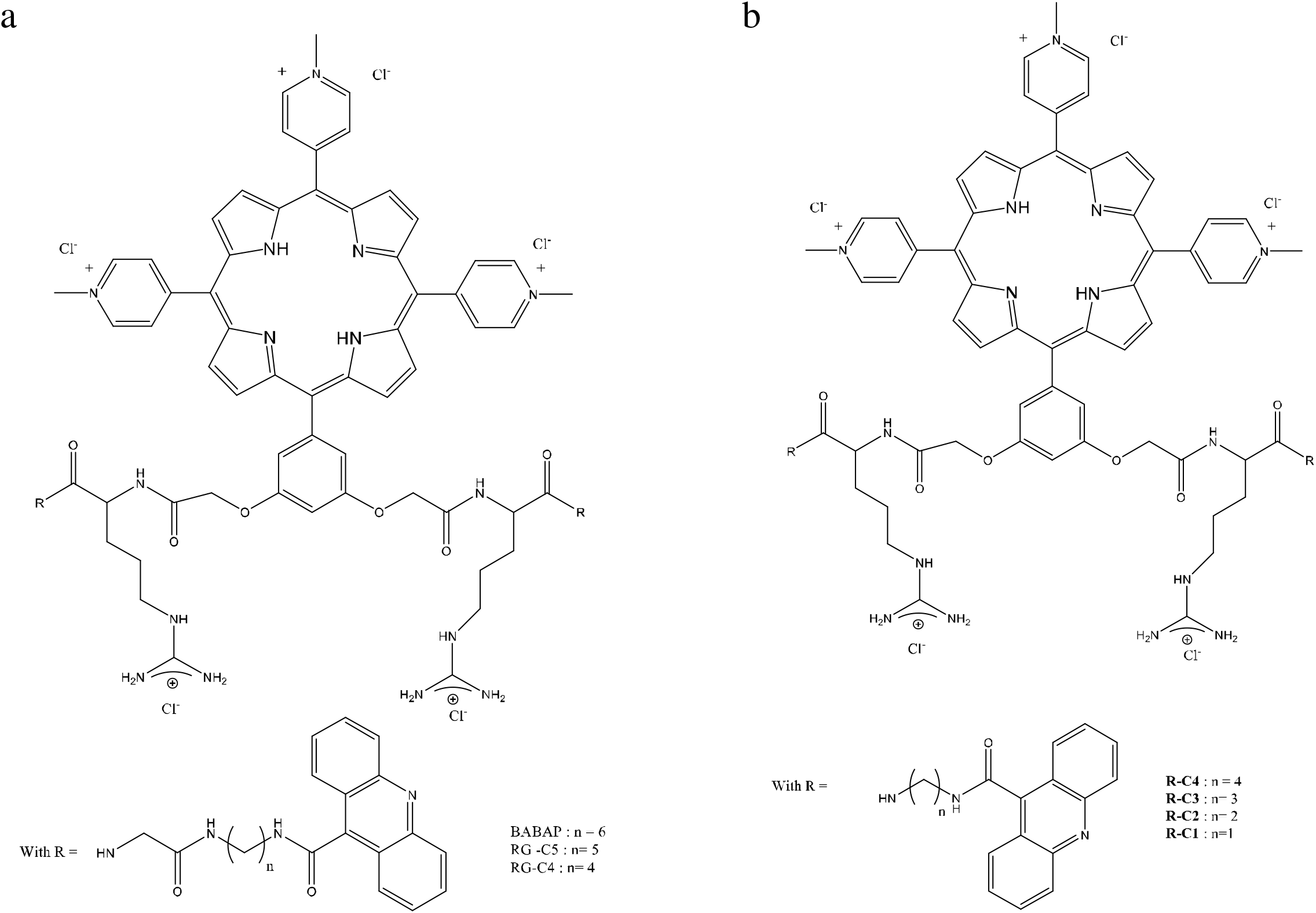
Representation of the molecular structures of BABAP, and its shorter derivatives RG-C5, RG-C4, and R-C4 till R-C1.

Kinked DNA conformations were reported earlier in complexes with proteins (19-25). This list is not exhaustive. It encompasses proteins such as CAP (21), (22), *Cre* recombinase (23), the CENP centromere protein (24), the TATA-binding box (25). The binding of intercalators and steroidal diamines was also reported to stabilize kinks. This was experimentally reported for ethidium bromide and actinomycin (26), irehdiamine (26), (27), (28), and, in the context of a JUMNA (29) energy-minimization study, for dipyrandium (30).

In the present study, the C6 chain connecting the Arg-Gly backbone is first shortened by one and two methylene groups, yielding the C5 and C4 analogs, respectively, of BABAP. This is followed by removal of the Gly backbone, and by up to three methylene groups, yielding the –C4 to -C1 Arg analogs of BABAP. The structural formulas of BABAP and its shorter-chain analogs are given in Figure 1. For each analog, we repeat an identical computational protocol, as detailed below. It consists of restrained energy-minimization then NVT equilibration, followed by limited-duration restrained MD and long-duration, unrestrained MD production. At its outcome, the most important structural features of the complexes and the complexed DNA conformation are analyzed.

### Computational procedure

We used the same procedure as in (14). MD simulations use the AMOEBA polarizable multipole potential (31) (and Refs. therein) coded in the Tinker-HP software (12) in its recent GPU implementation (13). We use the 2018 DNA parameters (31). The distributed multipoles of all derivatives compounds were derived by a Stone analysis (32) on their molecular wave functions computed with the B3LYP DFT functional (33) (34) and a cc-pVDZ basis set (35) (36) with the G09 (37) software.

All simulations were done on the 18-base palindromic sequence d(CGTA**C G**GG**C G**CC**C G**TACG)_2_, the same sequence as in the complex of a trisintercalating ligand, denoted **III** in (14). **III** is an oligopeptide derivative of an anthraquinone drug, mitoxantrone, the two arms of which were extended with an oligopeptide motif with an alkydiammonium side-chain connected by an amide-imidazole motif to an AA ring (14).

The starting trisintercalated DNA conformation was extracted from the previous MD simulation of its complex with ligand **III**. In a first step, manual docking was performed with the help of computer graphics with the Insight II software (Accelrys Inc., San Diego). The porphyrin ring was intercalated in the central d(CpG)_2_ sequence, locating its three cationic N-methylpyridinium rings in the minor groove, and the phenyl ring in the major groove. This is the ring location corresponding to the lowest energy minima of the complexes of BABAP (11) and BAP (38) with oligonucleotides having the target central palindromic sequence. On each DNA strand, the guanidinium side-chain was oriented in order to face the two successive G bases upstream from the central intercalation site, subsequently denoted as G_4_-G_3_/G_4’_-G_3’_. Appropriate torsional rotations around the C-C and C-N bonds of the connector then enabled AA intercalation in the corresponding d(C_1_pG_2_).d(C_9_pG_10_) or d(C_1’_pG_2’_).d(C_9’_pG_10’_) intercalation site of the target ten base-pair sequence. Manual docking was completed with a first round of constrained energy-minimization, bearing on the sole ligand in the presence of the following constraints: for each Arg side-chain, a 2.0 Å distance between the imino proton and O_6_ of G_3_/G_3’_ and that of the *cis*-amino proton and O_6_ of G_4_/G_4’_; for each AA ring, a distance of 3.8 Å between its N atom and the N_3_ atom of the two G bases and the two C bases of the corresponding intercalation site. Intramolecular restraints were added. To retain planarity of each AA ring, two distance restraints of 7.3 Å were set between the two outermost C atoms, namely between atoms C_2_ and C_7_ and atoms C_3_ and C_8_. The planarity of all amide bonds were enforced by distance restraints of 3.15 Å between the carbonyl O and the amide H atom.

The DNA-ligand complex was then immersed in a bath of water molecules in a box of dimensions 51, 54 and 90 Å on the x, y, and z dimensions. Neutrality was ensured by placing 29 Na^+^ replacing the last 29 water molecules of the water bath. Periodic boundary conditions (PBC) were applied along with Smooth Particle Mesh Ewald (PME) (39). We used cutoff values of 12 Å and 9 Å for van der Waals and Ewald interactions, respectively.

We then followed a similar procedure to that of (14).

Restrained energy-minimization was resumed, now also relaxing DNA, water and counterions. Molecular Dynamics was subsequently run. Equilibration was started by 50 K stepwise raises in temperature for a duration of 1 ns at constant volume, from 0 to 300 K. Production was then started at 300 K with a Bussi thermostat (40) and at constant pressure 1 atm using a Monte-Carlo barostat. Coordinates were saved every 100 ps.

We have retained the same inter- and intramolecular constraints as the preliminary stage. Additional restraints were added in the NVT and first 48ns NPT stages, destined to preserve the H-bonds between the complementary WC bases at, and capping, the three intercalation sites. These were set to prevent premature distortions of these base pairs. In addition, as motivated in (14), six additional restraints were introduced, which prevent the ‘fraying’ of the two end GC base-pairs in the course of MD, namely 2.0 Å between the corresponding O_6_-HN_4_, N_1_H-N_3_, and N_2_H-O_2_ atoms. This inclusion has precedents in DNA MD simulations including those with polarizable potentials (31). This was justified by the limited length of the oligonucleotide. It prevents unwanted ‘danglings’ of the end base-pairs.

During the first 48 ns production stage, the ligand-DNA distance restraints were maintained, as well as those between the above-mentioned complementary H-bonded bases. This was motivated in Ref. (14): large amplitude motion of DNA and the solvent were observed during the initial phases of the production runs, which cause an excess kinetic energy transfer to the bound ligand, which could destabilize and disrupt its complex with DNA, as well as distort the H-bond patterns of the bases. Unrestrained MD was resumed past 48ns, retaining, however, the intramolecular restraints enforcing acridine, guanidinium and formamide planarity and the six restraints enforcing H-bonding between the two-end base-pairs. The length of such unrestrained production runs was in the 150-200 ns range.

Structural analyses of the DNA at the outcome of MD were done with the CURVES+ software (41). The intermolecular interaction energies between the four bases, C_1_, G_2_, C_5’_, G_6’_ making up one acridine intercalation site, d(C_1_G_2_).d(C_5’_G_6’_), and the acridine ring were computed by DFT quantum chemistry and decomposed into their five contributions: Coulomb (E_C_), short-range repulsion (E_X_), polarization (E_pol_), charge-transfer (E_ct_) and dispersion (E_disp_). This was done by the ALMOEDA procedure (42) coded in the QChem package (43). We resorted to the ωB97-XD DFT functional (44) and the cc-pVTZ(-f) basis set (36).

## Results and discussion

### BABAP

A representation of the last MD pose of the complex of BABAP with the target oligonucleotide is given in Figures 2a-c. Figure 2a gives an overall representation of the complex. Figures 2b-c show the detailed H-bond interactions which take place on each strand between the imino and *cis-*amino protons of the Arg side-chain and O_6_/N_7_ of the two targeted bases, G_4_-G_3_/G_4’_-G_3’_ as well as the H-bond between the Arg main-chain NH and G_6_/G_6’_ of the central intercalation site. Figures 3a and 3b show the (normalized) distribution functions of the H-bond distances between the first and second Arg side-chain, respectively, and the two targeted G bases. Figure 3c shows the corresponding distributions for the H-bond between each Arg main-chain NH group and O_6_ of bases G_6_/G_6’_ of the central intercalation site. Figures 4a-c show the corresponding time evolutions of these distances. The structural results are fully consistent with the energy-minimization reported earlier (11). Each Arg side-chain can align along the z axis enabling it to bind to the two successive G bases on the same strand. This mode of binding, rather than the more frequent one in which an Arg side-chain binds coplanar to a single G base in a bidentate fashion, was reported in an X-ray study of the complex between yeast RAP1 protein and telomeric DNA. Thus Arg_404_ was found to bind to N_7_ of two successive G bases, G_8’_ and G_9’_ on the same strand (45). In the present complex, however, the interaction takes place preferentially with O_6_. With both arms, the distance distributions are much better peaked around O_6_ than around N_7_ (figures 3a-b). The distributions around O_6_ of G_6_/G_6’_ (figure 3c) are shallower, possibly due to the neutrality of the main-chain Arg NH group, compared to the partly positively charged imino and amino side-chain groups.

**Figures 2a-c.**
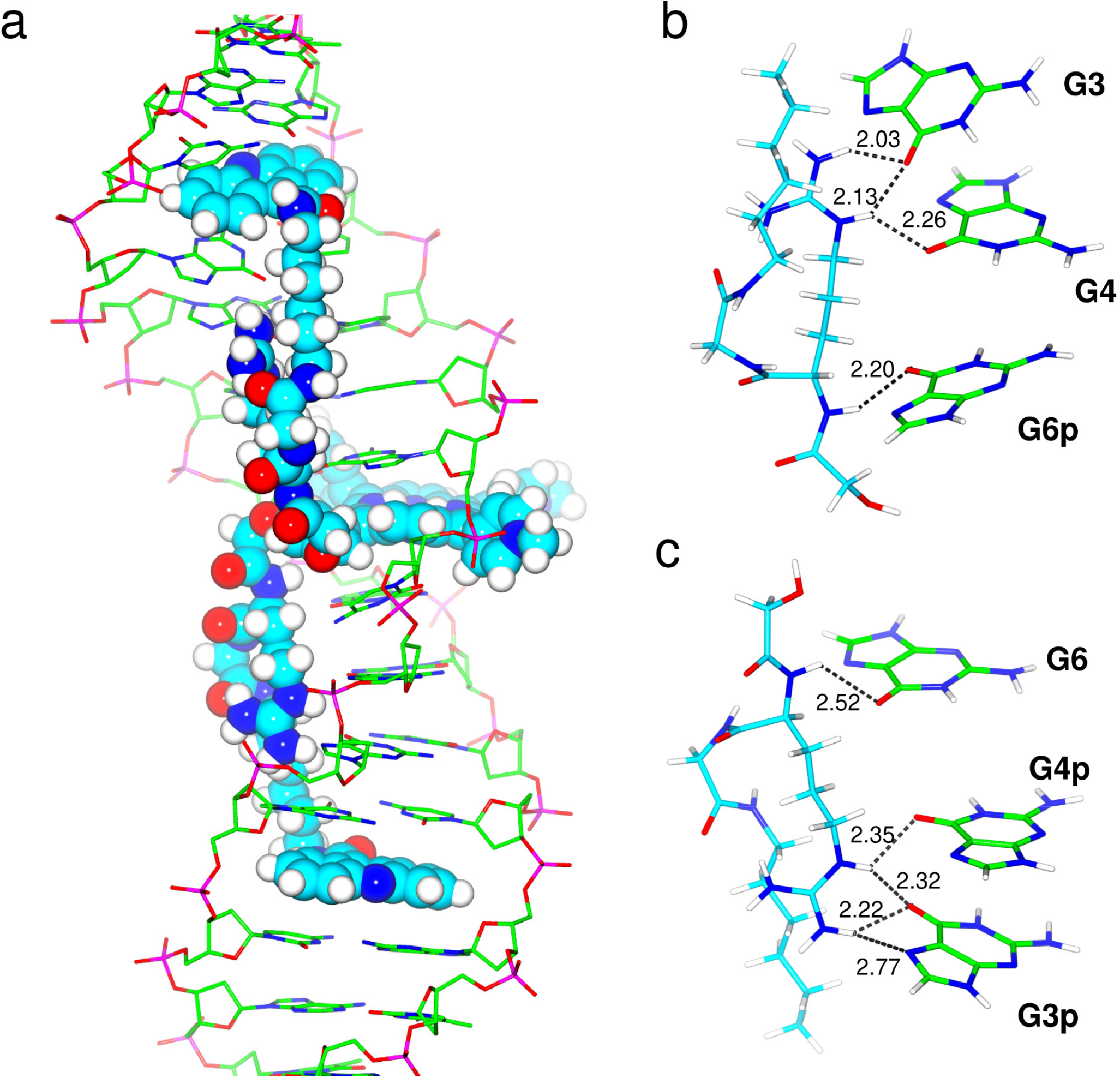
a) Overall representation of the DNA complex of BABAP. b-c) detailed H-bond interactions on the unprimed (b) and primed (c) strands between the imino and *cis-*amino protons of the Arg side-chain and O_6_/N_7_ of G3-G4/G3’-G4’.

**Figures 3a-c.**
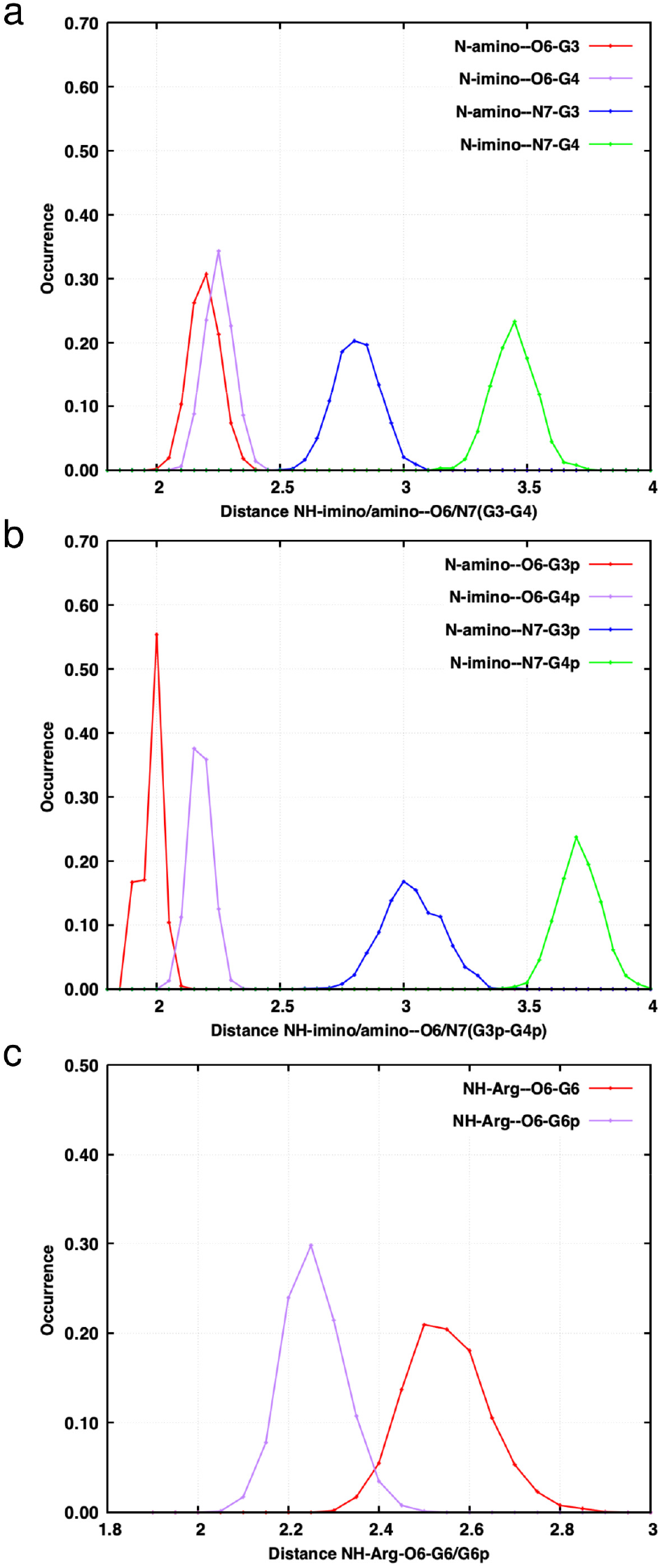
Normalized distribution functions of the H-bond distances between each Arg side-chain and the bases it targets in the unprimed (2a) and primed (2b) strands. 3c. Normalized distributions for the H-bond between each Arg main-chain NH group and O_6_ of bases G_6_/G_6’_ of the central intercalation site.

**Figures 4a-d.**
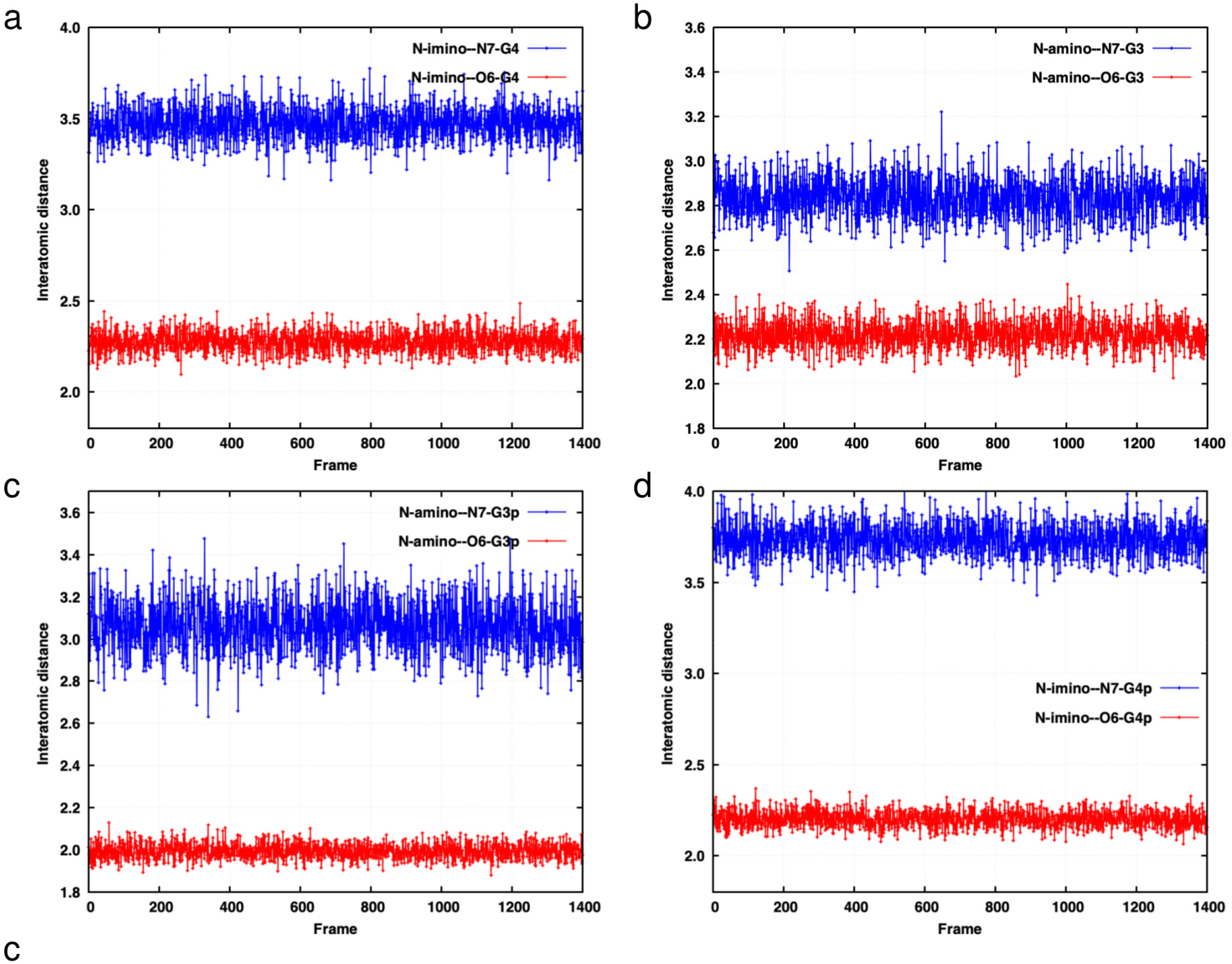
Time evolutions of the H-bond distances represented in 3a-c.

### Shorter BABAP derivatives

#### 1) Arg-Gly with a C5 or a C4 connector

Using the one-letter code for the peptide backbone, these derivatives will be denoted RG-C5 and RG-C4. We retain the acronym BABAP for the initial derivative rather than RG-C6.

*RG-C5*. An overall view is given in Supp. Info S1a, while the H-bond distances with G_3_-G_4_/G_3’_-G_4’_ are represented in Supp. Info S1b-c.

*RG-C4*. An overall view is given in Supp. Info S2a, while the H-bond distances with G_3_-G_4_/G_3’_-G_4’_ are represented in Supp. Info S2b-c.

Figure 5 represents the structure of DNA bound by BABAP, along with those with RG-C5 and RG-C4. It displays the distance between one P atom, P_8_, of the central intercalation site, and the two P atoms, P_14_ and P_31_, of the d(TpA)_2_ step six base-pairs downstream. P_8_ is closer to P_31_ on the opposite strand by 4.9 Å in the BABAP complex. This difference is comparable to those found with RG-C5, namely 4.1 Å and it increases to 5.7 Å with RG-C4.

**Figure 5.**
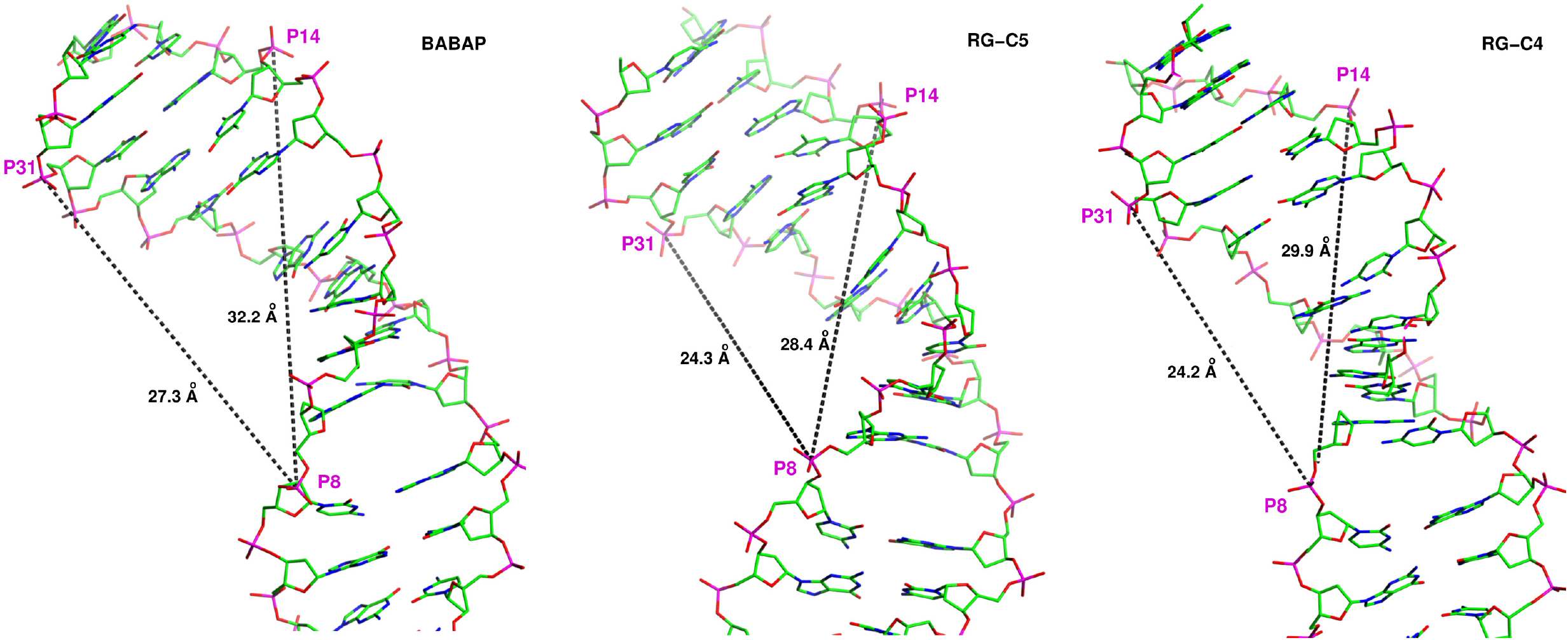
Representation of the structure of DNA bound by BABAP, alongside with those in its complexes with RG-C5 and RG-C4.

We next monitor the impact of further shortenings of the connecting chain, which follow the removal of the Gly backbone.

#### 2) Arg with C4 to C1 connectors

With the one-letter code, these derivatives will be denoted as R-C4 till R-C1.

Representations of the overall complexes of R-C4 till R-C2, and of the Arg H-bonds with the G bases are given as Supp. Info S3-S5. Figures 6a-d represent the structures of DNA in its complexes with the four derivatives focusing on the P_8_-P_14_ and P_8_-P_31_ distances. Figure 7a gives a representation of the kinked DNA---R-C1 complex and figures 7b-c display the relevant H-bonds between the G bases and Arg from the last MD pose. On the unprimed strand, the interactions of the Arg side-chain involve only the amino hydrogens. Each N atom donates its H atom *trans* to the imino proton to N_7_ of G_4_, and both amino hydrogens belonging to the N atom *cis* to the imino proton are donated to N_7_ to G_3_. Owing to the ‘pinching’ of the major groove, the amino H *cis* to the imino proton is also donated to N_7_ of G_2_, one more step upstream, and on the 5’site of the AA intercalation site. The carbonyl O of the peptide bond preceding the Arg residue accepts a proton from the extracyclic amino group of cytosine C_5_ on the 3’side of the porphyrin intercalation site. On the primed strand, the imino proton and its *cis*-amino proton bind to G_4’_ and G_3’_, respectively but with N_7_ rather than with O_6_ as was the case with BABAP. The main-chain NH is donated to N_7_ of G_6_ of the intercalation site on the unprimed strand.

**Figures 6a-d.**
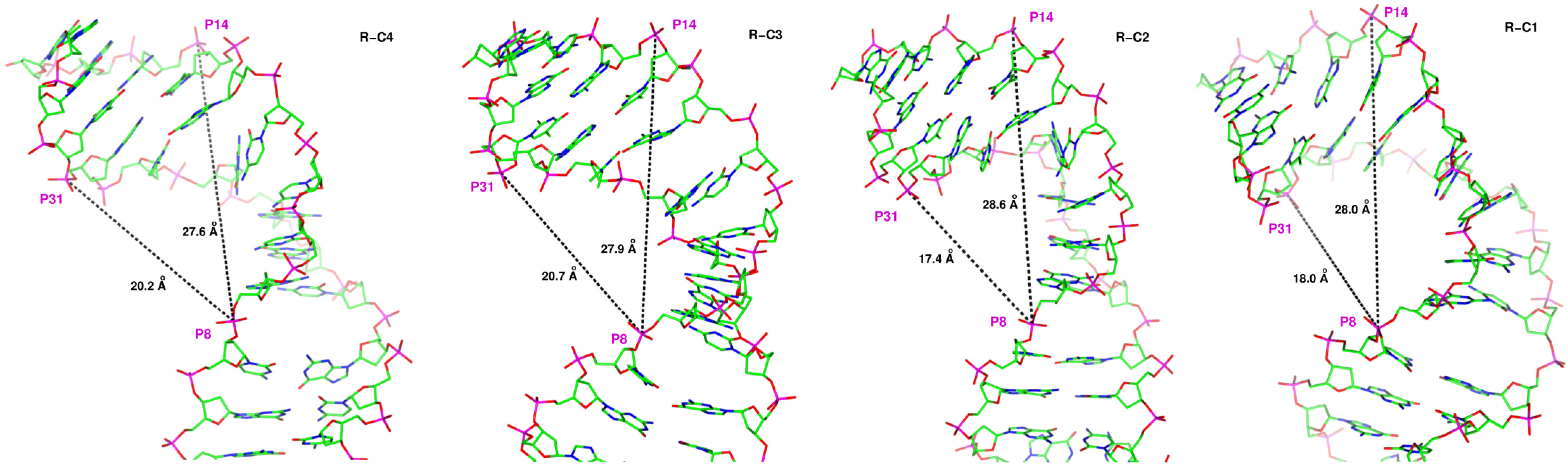
Representation of the structures of DNA in its complexes with derivatives R-C4(a), R-C3(b), R-C2(c), and R-C1(d) displaying the P_8_-P_14_ and P_8_-P_31_ distances (see text for definition).

**Figures 7a-c.**
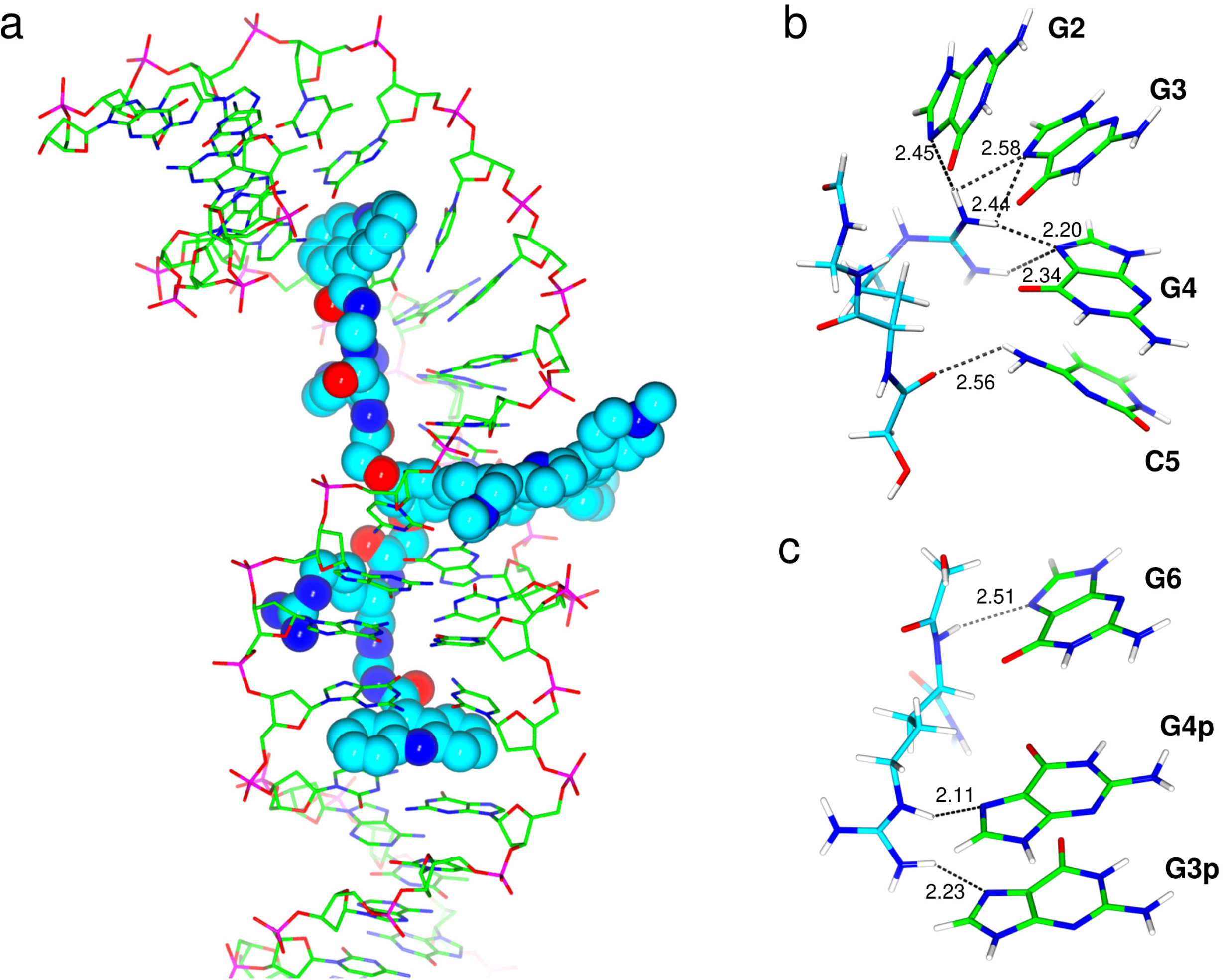
a) Representation of the kinked DNA---R-C1 complex. b-c) Representation of the H-bonds between the G bases and Arg from the last MD pose, on the unprimed (b) and primed (c) strands.

Figures 8a-b give the normalized distributions of the relevant Arg-guanine H-bond distances. On the unprimed strand, three of the five H-bonds represented in Figure 7a have their peaks at 2.2-2.3 Å. The first H-bond involving the proton *cis* to the imino proton (denoted as N_c_H_c_) and N_7_ of G_3_, has a slightly shifted peak at 2.35 Å, while the second H-bond, donated to N_7_ of G_2_, is shifted to 2.6 Å. On the primed strand, the distributions are much more accented with marked peaks at similar distances, 2.1 and 2.2 Å for the imino and *cis*-amino hydrogens, respectively. The corresponding structural informations of the DNA complexes with R-C4 till R-C2 are given as Supp. Infos 3-6.

**Figures 8a-b.**
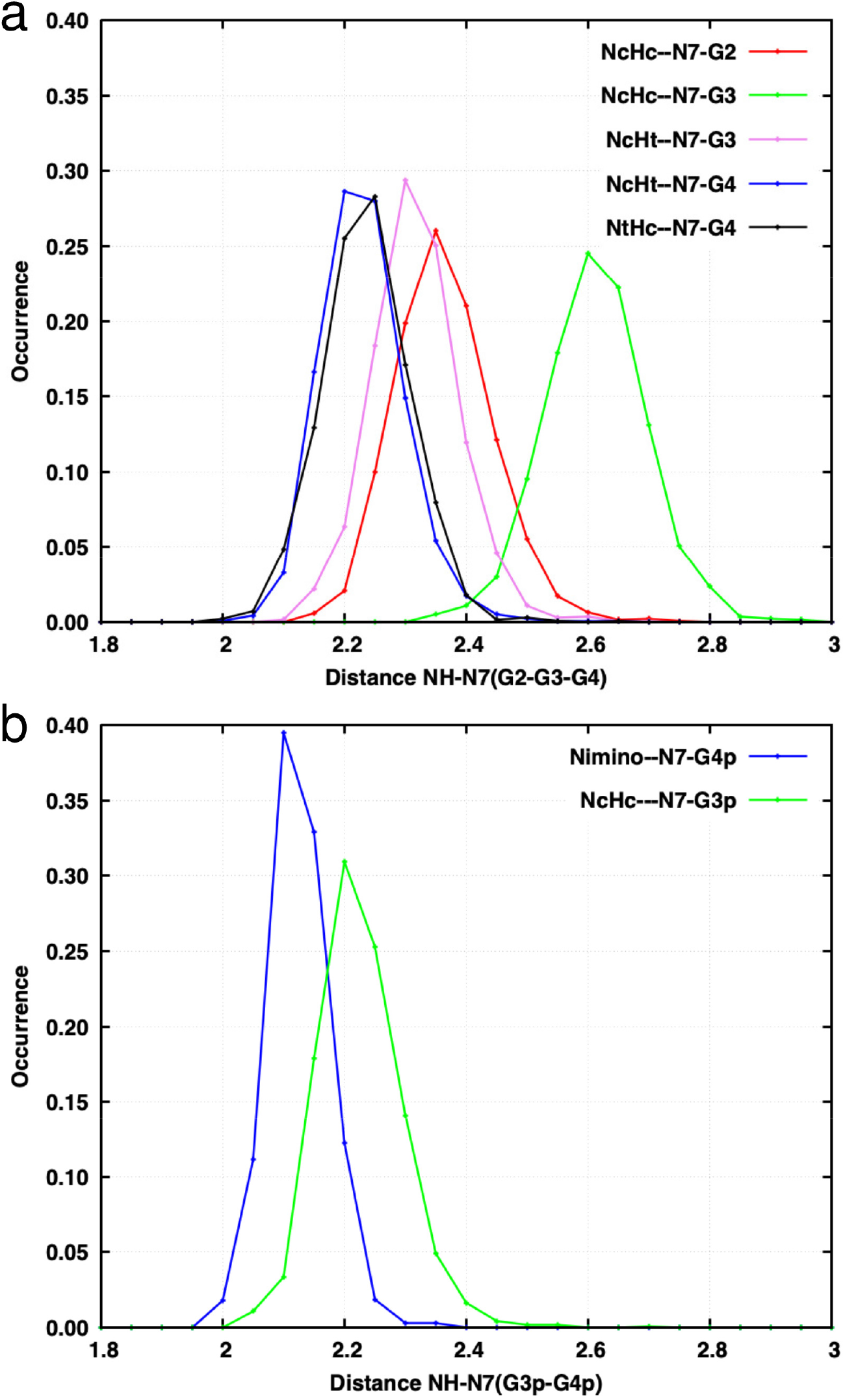
DNA--R-C1 complex. Normalized distributions of the Arg-guanine H-bond distances on the unprimed (a) and primed (b) strands.

A revealing indicator of the progressive ligand-induced DNA kink is the P_8_-P_31_ distance. It decreases from 27.3 Å in the BABAP complex to 24.3 Å in those with RG-C5 and RG-C4, then to 20.7-20.5 Å with R-C4 and R-C3, and has its smallest values, 17.4-18.0 Å, with R-C2 and R-C1. The difference between the P_8_-P_14_ and P_8_-P_31_ distances has increased between the two ‘extreme’ DNA ligands, BABAP and R-C1, from 4.9 to 10 Å. Relevant P-P distances are reported in Table I.

**Table I.** Values (Å) of the across-strand P8-P31 and along-strand P8-P14 distances and values of their differences in the DNA complexes with BABAP and derivatives with progressive linker shortenings (see text for definition).

| Ligand | P8-P31 | P8-P14 | Difference<br>(P8-P14)-(P8-P31) |
| --- | --- | --- | --- |
| BABAP (RG-C6) | 27.3 | 32.2 | 4.9 |
| RG-C5 | 24.3 | 28.4 | 4.1 |
| RG-C4 | 24.2 | 29.9 | 5.7 |
| R-C4 | 20.2 | 27.6 | 7.4 |
| R-C3 | 20.7 | 28.1 | 7.4 |
| R-C2 | 17.4 | 28.6 | 11.2 |
| R-C1 | 18.0 | 28.0 | 10.0 |

Figure 9 recasts the structures of DNA in its R-C1 complex, along with its structure extracted from its complex with the *Cre* recombinase (23). The R-C1 complex bears a striking resemblance with the kinked *Cre*-bound segment of DNA. The two P-P distances corresponding to the P_8_-P_14_ and P_8_-P_31_ ones amount to 15.6 and 26.1 Å, closely comparable to the respective ones of 18.0 and 28.0 Å in R-C1. Such structural similarities imply that the R-C1 induced kinks are experimentally accessible. It could also be noted that the bases at the level at the kink site (denoted as ‘the second and third base pairs from one end of the central crossover region’ in Guo et al. 1999) have a negative roll of −49° and a positive tilt of 16°. The roll and tilt values found for the R-C1 complex at the AA intercalation site, as determined by the Curves+ software, are −72° and 33°, respectively. Such negative roll and positive tilt values could not, however, be generalized to the other complexes from the present study, nor in the other experimental DNA-protein complexes from Refs. (20-22, 24-25), which have less accented DNA kinks. For the last pose of the BABAP and of the R-C1 DNA complexes, we have computed the intermolecular interaction energies stabilizing the stacking of the acridine ring with the four bases, C_1_, G_2_, C_5’_ and G_6’_ of the d(C_1_G_2_).(C_5’_G_6’_) dinucleotide. The acridine ring had its amide group substituted with a terminal -CH_3_ group. The four bases were disconnected from their sugars and the valence of their N atoms, N_1_ or N_9_, completed by an H atom replacing the C_1’_ sugar. ΔE_tot_ embodies the acridine-base interactions and the base-base interactions. The results of the ALMOEDA energy analyses are reported in Table II. ΔE_tot_ has close values in the BABAP and R-C1 complexes, and the individual energy contributions have comparable magnitudes in both. This indicates that the stacking of the acridine is not impaired by the distortion of DNA that occurs in the R-C1 complex.

**Figure 9.**
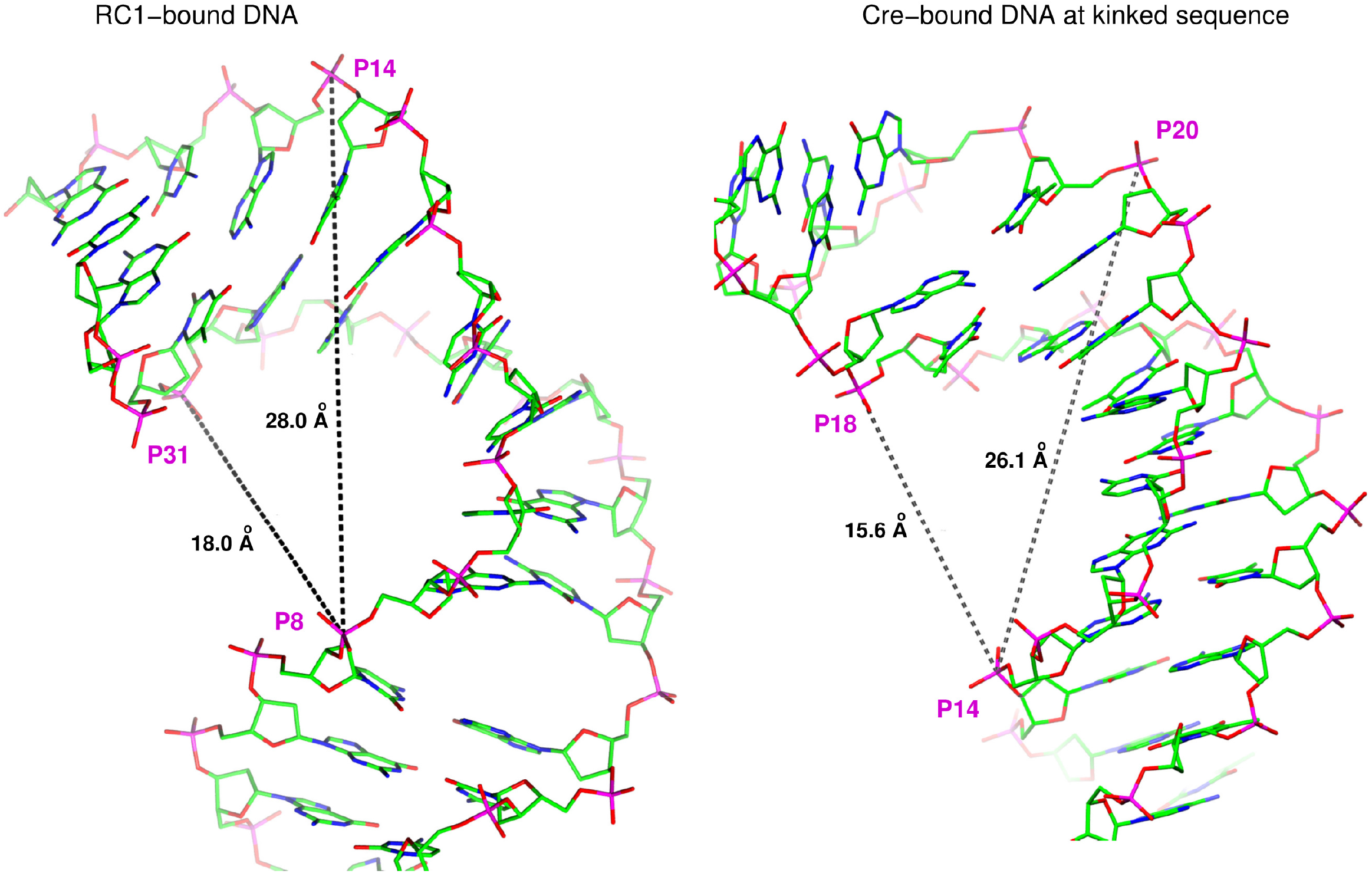
Structure of DNA in its R-C1 complex, alongside with its structure extracted from its complex with the *Cre* recombinase.

**Table II.**
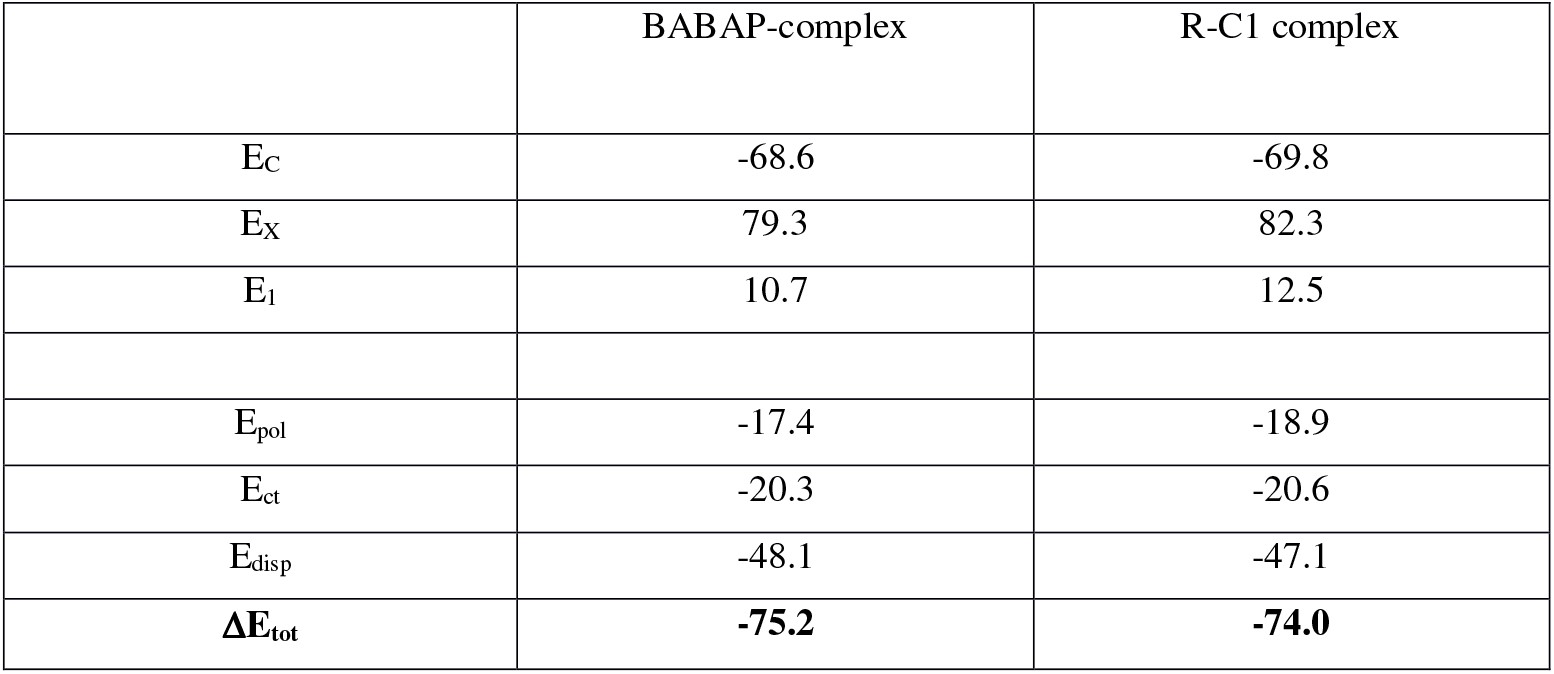
Intermolecular interaction energies in the stacked complex of the acridine ring with the four bases, C_1,_ G_2_, C_5’,_ G_6’_, of the d(C_1_G_2_).d(C_5’_G_6’_) sequence extracted from the complex of BABAP and R-C1 ligands with the target 18-mer in the last MD pose. See text for definition. Energies in kcal/mol.

### Conclusions and Perspectives

The present polarizable MD simulations show the possibility of inducing sequence-selective, progressive deformations on DNA along a series of its complexes with trisintercalating oligopeptide derivatives. In this study, the starting ligand is BABAP (bisacridinyl-bisarginyl porphyrin). It has a tricationic porphyrin which intercalates at the central d(CpG)_2_ sequence of the d(C_1_G_2_G_3_G_4_C_5_G_6_C_7_C_8_C_9_G_10_)_2_ palindrome. The phenyl ring is substituted by an Arg-Gly (RG) peptide connected by a C6 linker to an AA ring which intercalates at an end d(CpG)_2_ sequence. Trisintercalation was previously demonstrated by circular dichroism and by Topoisomerase I unwinding experiments (11). MD showed the persistence of the H-bonds between the imino proton of each Arg side-chain and O_6_ of G_4_/G_4’_ and of its *cis*-amino proton with O_6_ of G_3_/G_3’_. There were no distortions of the sugar-phosphate backbone along the non-intercalated sequences. Progressive shortenings of the linker chain to the AA ring were subsequently performed, from C6 to C4, followed by removal of the Gly backbone and further shortenings, from C4 till C1. MD simulations on the six DNA-ligand complexes showed DNA to undergo distortions (‘kinks’) of increasing magnitude. A sensitive indicator is the distance between one P atom of the central intercalation site, denoted as P_8_, and, on the opposite strand, the P atom (P_31_) of the d(TpA)_2_ dinucleotide six base-pairs downstream. It underwent decreases from 27.3 Å in the BABAP complex, to 24.3 Å in the RG-C5 and RG-C4 complexes, to app. 20.5 Å in the R-C4 and R-C3 ones, to 17.4-18.0 Å in the R-C2 and R-C1 complexes.

The kinked structure of the DNA-R-C1 complex bears a striking resemblance to that of a DNA complexed with *Cre* recombinase. It is highlighted in Figure 9. This proves that such ‘extreme’ kinked structures could be energetically accessible to DNA. Small DNA-ligands are also able to kink DNA. Early examples are provided by steroid diamines (26) (27) (28) and the bisintercalator ditercalinium (46) (47), even though the DNA deformation (48) is less than with the extreme compounds R-C2 and R-C1. Kinked DNA structures are hypersensitive sites for cleavage by endonucleases (49). This implies that, along with their DNA photosensitizer properties, some BABAP derivatives could also, in the presence of endonucleases, act to induce sequence-selective DNA cleavage.

Induction of kinks of modulable amplitudes could, alternatively, be enabled by replacing one, or both, AA intercalator(s), by groove binders having varying lengths and appropriate shape with polar groups destined to bind O_6_/N_7_ of G_2_/G_1’_, and/or bind to the extracyclic –NH_2_ of C_1_/C_5’_ (Gresh et al., work in progress). Conversely, intercalation into the d(C_1_-G_2_).d(C_5’_G_6’_) sequence could be enhanced with conjugated rings with augmented stacking affinities for it. This could be achieved with intercalators having enlarged rings, the kinks being enforced, as with AA, by controlled connector shortening. One possibility currently investigated is AA replacement by a Ru(II) polypyridyl complex of Ru(II) (50). Induction of kinked DNA was also found by us in a novel series of anthraquinone derivatives related to compound **III** of (14) and will be reported in due course.

Along with sequence-selectivity and binding affinities, novel DNA-binding oligopeptide derivatives of intercalators could thus be tailored in order to *de novo* modulate and control the amplitude of the kink they induce in the course of dynamics. An initial docking at a predefined kinked DNA conformation of related compounds is first done by energy-minimization. The structures of the complexes are then evolved in the course of MD, leading to the selection of those derivatives with connectors enforcing and maintaining the kinked conformation over the duration of MD. Accordingly, the structures of the DNA complexes with BABAP and R-C1 are provided as Supp. Info. These should enable such simulations on related derivatives having a central intercalating ring from other families, such as anthraquinone, anthracycline, or aminoacridine. Additional data are available upon request.

## Supporting Information

**Figures S1.** Complex of DNA with RG-C5. Overall view (S1a) and H-bond with G_3_-G_4_ (S1b) and G_3’_-G_4’_ (S1c).

**Figures S2.** Complex of DNA with RG-C4. Overall view (S2a) and H-bond with G_3_-G_4_ (S2b) and G_3’_-G_4’_ (S2c).

**Figures S3.** Complex of DNA with R-C4. Overall view (S3a) and H-bonds in the unprimed (S3b) and primed (S3c) strands.

**Figures S4.** Complex of DNA with R-C3. Overall view (S4a) and H-bonds in the unprimed (S4b) and primed (S4c) strands.

**Figures S5.** Complex of DNA with R-C2. Overall view (S5a) and H-bonds in the unprimed (S5b) and primed (S5c) strands.

## Data and Software Availability

Four AMOEBA MD input files are provided for BABAP and ligand R-C1, with extensions *xyz (coordinates), *key and *prm (parameters) and *dyn (for MD restarts) and regrouped for each ligand as one file (lig-*-MD.pdf). They are provided with the paper as data files. The calculations were performed with the publicly available free open-source software Tinker-HP (https://github.com/TinkerTools/tinker-hp).

## Acknowledgements

We wish to thank the Grand Equipement de Calcul Intensif (GENCI): Institut du Développement et des Ressources en Informatique (IDRIS), Centre Informatique de l’Enseignement Supérieur (CINES), France, project x-2009-07509, and the Centre Régional Informatique et d’Applications Numériques de Normandie (CRIANN), project 19980853.

---

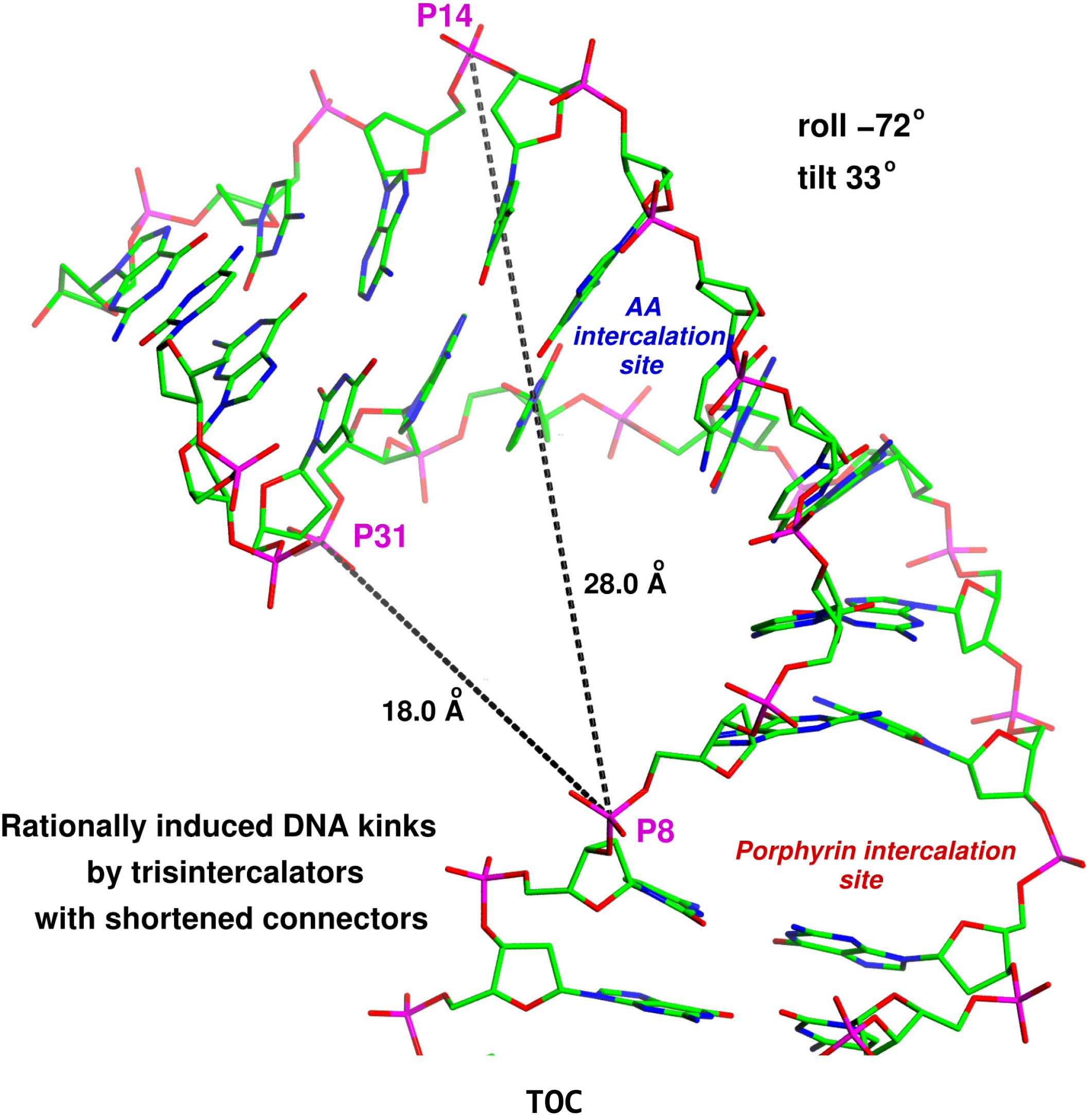

## Notes

### Competing Interest Statement

The authors have declared no competing interest.

